# Characterizing neural coding performance for populations of sensory neurons: comparing a weighted spike distance metrics to other analytical methods

**DOI:** 10.1101/778514

**Authors:** G. Marsat, K.C. Daly, J.A. Drew

## Abstract

The identity of sensory stimuli is encoded in the spatio-temporal patterns of responses of the encoding neural population. For stimuli to be discriminated reliably, differences in population responses must be accurately decoded by downstream networks. Several methods to compare patterns of responses have been used by neurophysiologists to characterize the accuracy of the sensory responses studied. Among the most widely used analysis, we note methods based on Euclidean distances or on spike metric distances. Methods based on artificial neural networks and machine learning that recognize and/or classify specific input patterns have also gained popularity. Here, we first compare these three strategies using datasets from three different model systems: the moth olfactory system, the electrosensory system of gymnotids, and leaky-integrate-and-fire (LIF) model responses. We show that the input-weighting procedure inherent to artificial neural networks allows the efficient extraction of information relevant to stimulus discrimination. To combine the convenience of methods such as spike metric distances but leverage the advantages of weighting the inputs, we propose a measure based on geometric distances where each dimension is weighted proportionally to how informative it is. We show that the result of this Weighted Euclidian Distance (WED) analysis performs as well or better than the artificial neural network we tested and outperforms the more traditional spike distance metrics. We applied information theoretic analysis to LIF responses and compare their encoding accuracy with the discrimination accuracy quantified through this WED analysis. We show a high degree of correlation between discrimination accuracy and information content and that our weighting procedure allowed the efficient use of information present to perform the discrimination task. We argue that our proposed measure provides the flexibility and ease of use sought by neurophysiologists while providing a more powerful way to extract relevant information than more traditional methods.

## INTRODUCTION

Encoding of sensory signals is typically mediated by the patterned spiking responses of a population of sensory neurons (Stanley, 2013). Various aspect of this spatio-temporal pattern can represent the identity information carried by the population response and the reliability of the response reflect the accuracy of encoding (Rieke et al., 1997). Several methods have been developed to characterize the encoding accuracy and better understand the coding strategy. Three commonly used methods are: 1-Analyses based on information theoretic calculation (e.g. Clague et al., 1997; Rieke et al., 1997); 2-Spike distances metrics (e.g. Victor, 2005; Allen and Marsat, 2019) often paired with ROC analysis and 3-Pattern classifiers based on artificial neural network (e.g. Barrett et al., 2019; Glaser et al., 2020). Our goal is to compare these methods and explore an alternative that combines the advantages of different approaches.

Information theory (Shannon, 1948) has been applied to neural systems leading to impressive insight into sensory processing (Bialek and Rieke, 1992). Direct methods to quantify the information content of neural responses about a set of stimuli typically require large datasets so alternative methods using white noise stimuli have been developed (Rieke et al., 1997). Although these methods have been applied extensively and provided many insights into neural coding (Passaglia and Troy, 2004; Marsat and Pollack, 2005; Middleton et al., 2009), the output measure quantifies coding accuracy (typically in bit/s) but it is hard to relate it to behavioral performance in response to natural stimuli. Spike distance metrics provide useful alternatives to information-theoretic approaches because they can be applied to datasets of reasonable size using naturalistic stimuli. These spike-distance metrics rely on quantifying the similarity between spike trains by either transforming one spike train into the other (with each step in the transformation being associated with a cost; Victor and Purpura, 1997); using the integral of the difference between spike trains that have been convolved with a smoothing kernel (van Rossum, 2001); or calculating the Euclidean distance of multidimensionally mapped neural responses (Daly et al., 2004b; Kreher et al., 2008). Many variations of these measures have been tested including versions that consider populations of neural responses (Houghton and Sen, 2008) of measures that rely on non-Euclidean measures (Wesolowski et al., 2014; Guo et al., 2022). These measures are convenient because they can easily be paired with ROC-type discrimination analysis and thus lead to a performance estimate that can be compared directly to behavioral performance (Parnas et al., 2013; Allen and Marsat, 2018). Furthermore, these decoders based on spike metric distances emulate a decoding process that is biologically realistic (Larson et al., 2010). It is not clear, however, how these types of decoders are optimized. Artificial neural networks (ANNs) such as the ones used in machine learning, leverage powerful algorithms to maximize the use of information-rich parts of the input and weigh down the noisy portions of the input. ANNs can learn to associate specific aspects of the neural responses with particular stimuli and thus can result in very efficient classifiers (Szabó and Barthó, 2022). Although this approach can be very successful when the goal is simply decoding the neural responses, it is not clear how the process relates to the performance of actual neural systems. The weights and the dynamic of the ANN are mostly hidden from the experimenter and it is unlikely that the different components of the decoder can be mapped onto a similar process in the biological system. This approach is thus limited for the purpose of understanding how sensory systems encode and decode information.

In this paper, we compare the performance of different methods and present a new approach that combines the convenience and biological realism of spike-metric decoders but leverages the input-weighting approaches that allow ANNs to be so efficient. To do so, we use three datasets that cover a broad range of neural coding scenarios in lower sensory systems. We use recordings from olfactory responses of projection neurons (PNs) in the antennal lobe (AL) of moth (Daly et al., 2016), pyramidal cell (PC) responses to communication stimuli in the electrosensory lateral line lobe (ELL) of weakly electric fish (Allen and Marsat, 2018) and responses of leaky integrate and fire (LIF) model neurons to frozen white noise in a linear regime. Each system encodes the relevant information in different aspects of the population response’s spatiotemporal pattern. We ask how accurately we can extract the relevant information from these responses and reliably discriminate between sensory stimuli. We show that simply adding an input-weighting procedure to a spike-metric distance decoder allows a decoding performance similar to-or even surpassing-a simple ANN while retaining a biologically-plausible process.

## METHODS

### Datasets

*In vivo* neural recordings were obtained from previous research and the details of the experiments are given in the corresponding publications (Daly et al., 2016; Allen and Marsat, 2018). Briefly, the olfactory responses consisted of one-second-long projection neuron (PN) recordings from the antennal lobe (AL) of *Manduca sexta* moths following odor presentation. Six different concentrated odors that are commonly used in this model system were used (100 ms long puffs) and one blank stimulus. The binarized spike trains were initially sampled at 2KHz, convolved with a 20 ms kernel (either a Gaussian function or an alpha function), and down-sampled to 100 Hz. Electrosensory responses consisted of responses of pyramidal cells (PCs) of the electrosensory lateral line lobe (ELL) when presented with communication signals (3 different type-1 chirps – “big chirps” – occurring on a high-frequency background beat). We used the responses of OFF cells, which encode the chirps better, and we analyzed a 350 ms window around the chirp timing (note that previously published results used a tighter 45 ms window which leads to more accurate discrimination results using an unweighted analysis). The binarized spike trains were sampled at 2 KHz and convolved with a 20 ms wide kernel (either a Gaussian function or an alpha function) to get a smooth estimate of the instantaneous firing rate.

A generic LIF model was used and parameters (threshold, capacitance, input current, noise strength) to obtain a firing rate modulation linearly correlated with the input. In a first version (figures 2–6) the threshold and capacitance of the neurons in the population were slightly varied by drawing from a normally distributed range of values. Since this heterogeneity could potentially lead to small non-linearities due to threshold and saturation, we used a homogeneous population for the comparisons with information theoretic measures in figures 7–8. Stimuli consisted of 1 second long, low-pass filtered, frozen white noise (0-40Hz), and 10 different patterns were used as the stimulus set. A population of 100 neurons was created with several repeats of responses to each stimulus; neural noise included in the LIF model was different from repeat to repeat. The neural noise was also different for each neuron, except for the analysis of noise correlations for which 1/3 of the total noise was shared across pairs of neurons (for a review on noise correlation and their significance, see: Kohn et al., 2016). Noise level and signal-to-noise ratios were varied in Figures 6–8 but were fixed at levels that lead to medium discrimination performance in previous figures. Responses were generated at a sampling rate of 20KHz, convolved with a gaussian or alpha function 20 ms wide, and down-sampled to 2 KHz.

**Figure 1:**
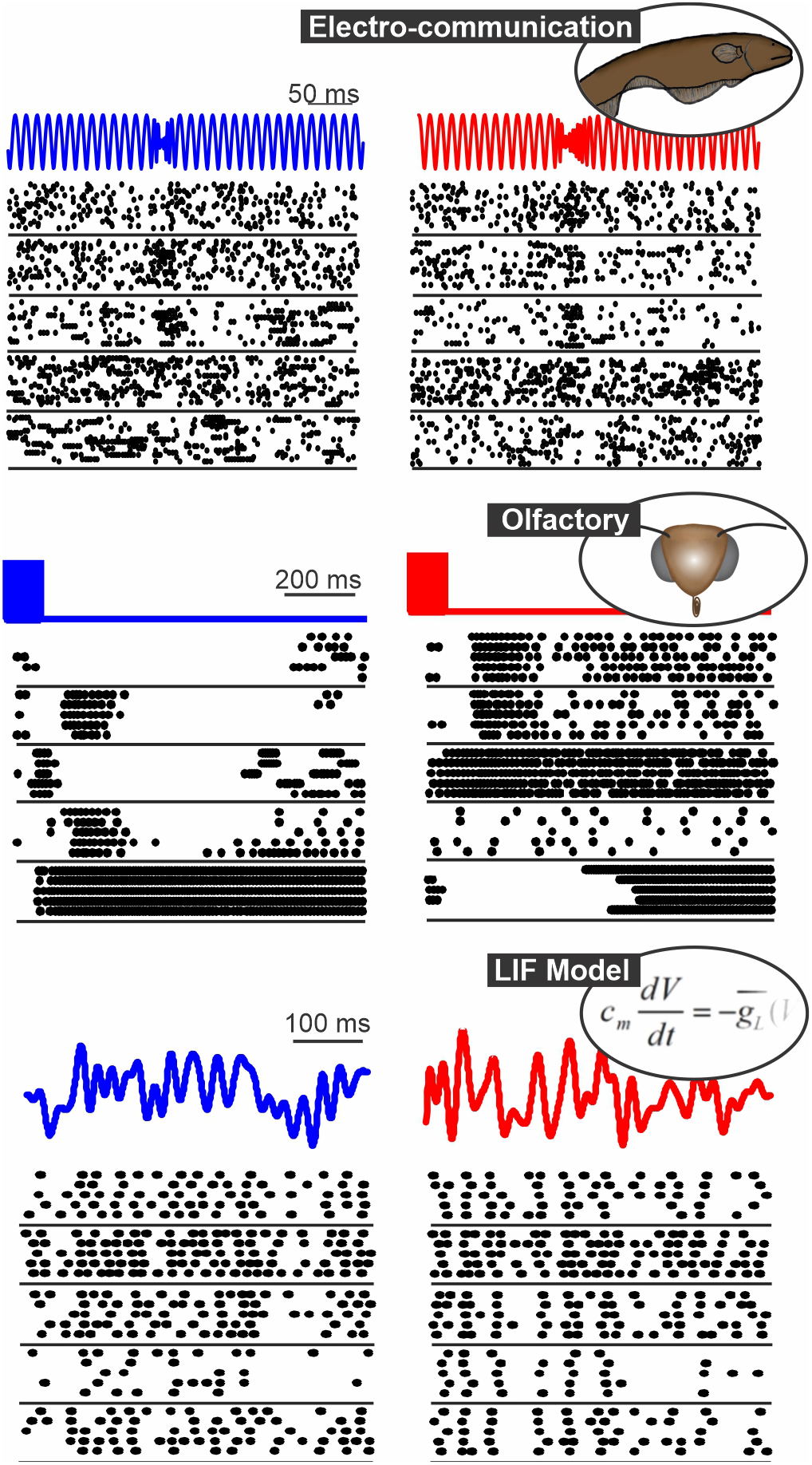
Spatio-temporal patterns of response in 3 systems: the gymnotid electrosensory system (responses of ELL pyramidal cells to chirps, Allen and Marsat, 2018), the moth olfactory system (PN responses to 2 odors, Daly et al., 2016) and a population of model neurons (LIF neurons stimulated with frozen white noise). In each panel, we show the responses of 5 neurons to repeated presentation of 2 different stimuli (blue vs red).

**Figure 2:**
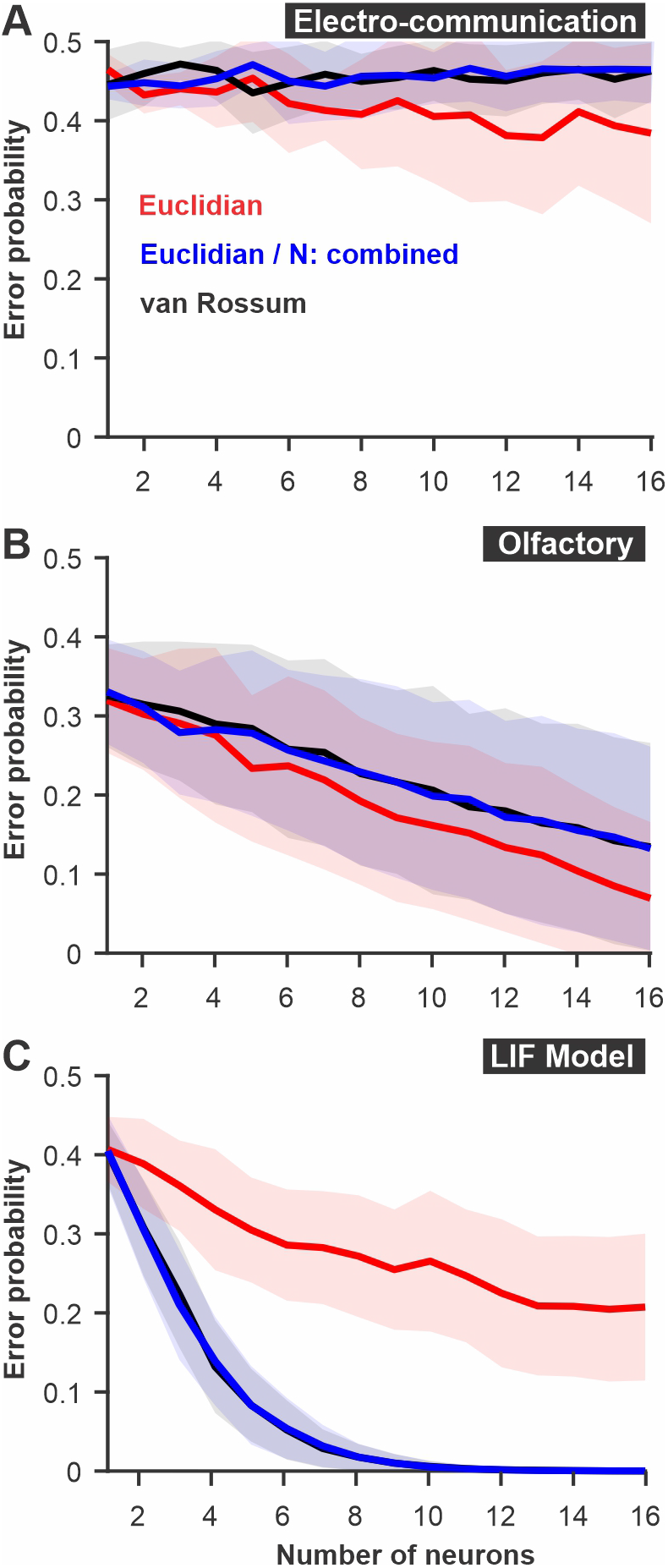
Discrimination accuracy for populations of neurons using established spike metrics. The probability of discrimination errors (y-axis), based on responses to pairs of different stimuli, was calculated using neural populations of varying sizes (x-axis). We compare methods based on Euclidean distance and the van Rossum spike distance metrics and show that they are very similar when the different neurons are not mapped as different dimensions in Euclidean space but combined (e.g. averaged into a population response). **A.** PC neuron responses to electrosensory chirps. **B**. PN response of the moth antennal lobe to odors. **C.** LIF model responses to frozen white noise stimuli of different shapes. Curves show averages (± s.d.) across all pairs of stimuli (number of stimuli: electrosensory=3 different chirps, 3 pairs; olfactory= 7 odors, 21 pairs; LIF=10 noise patterns, 45 pairs).

**Figure 3:**
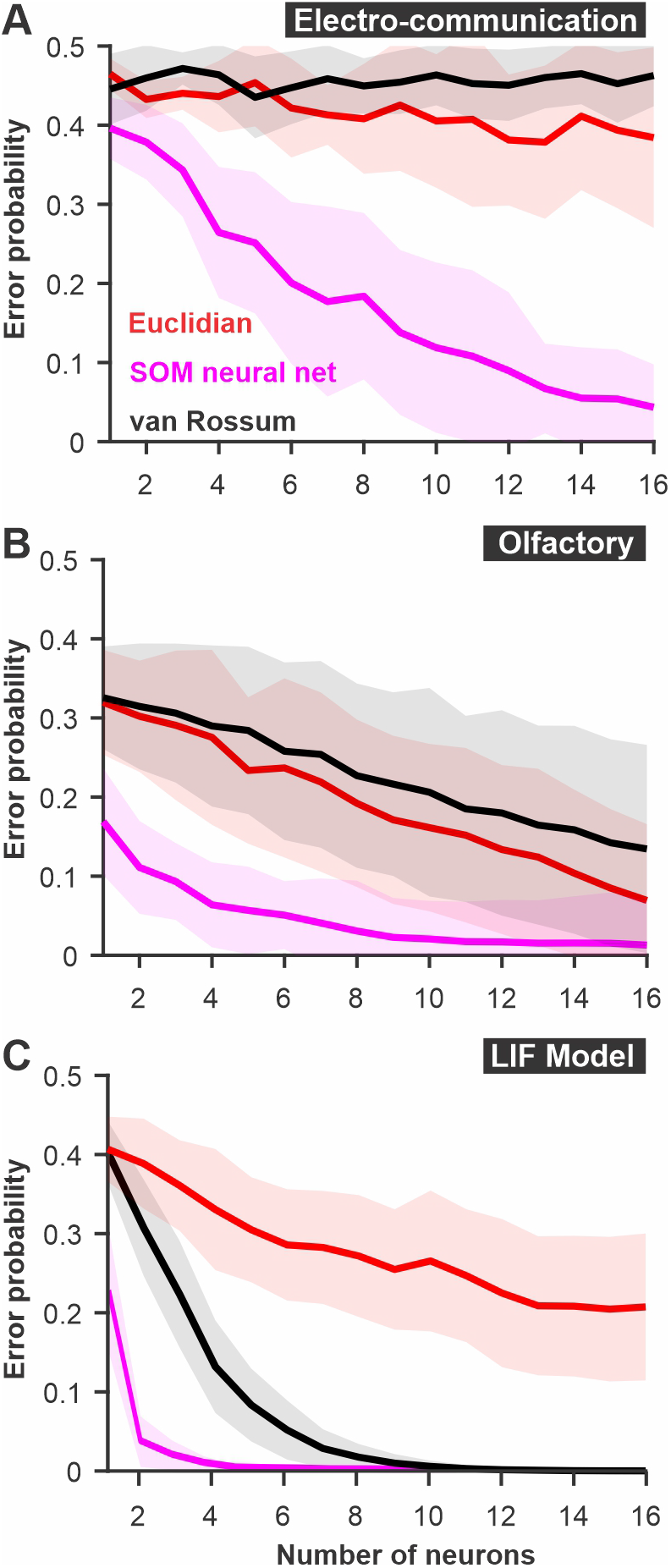
The discriminability of responses estimated using a SOM neural net. The spatio-temporally patterned array of inputs is weighted based on unsupervised learning to cluster the sets of inputs according to the variability and the patterns present in the dataset. Large intrinsic differences in patterns between responses to 2 stimuli thus lead to reliable clustering. We compare this SOM decoder where each timed point and each neuron are weighted independently (magenta) to a decoder based on Euclidean distance with time points and neurons kept as separate dimensions or a van Rossum metric where responses of different neurons are averaged together before the comparison. **A.** PC neuron responses to electrosensory chirps. **B**. PN response of the moth antennal lobe to odors. **C.** LIF model responses to frozen white noise stimuli of different shapes. Curves show averages (± s.d.) across all pairs of stimuli (number of stimuli: electrosensory=3 different chirps, 3 pairs; olfactory= 7 odors, 21 pairs; LIF=10 noise patterns, 45 pairs).

**Figure 4:**
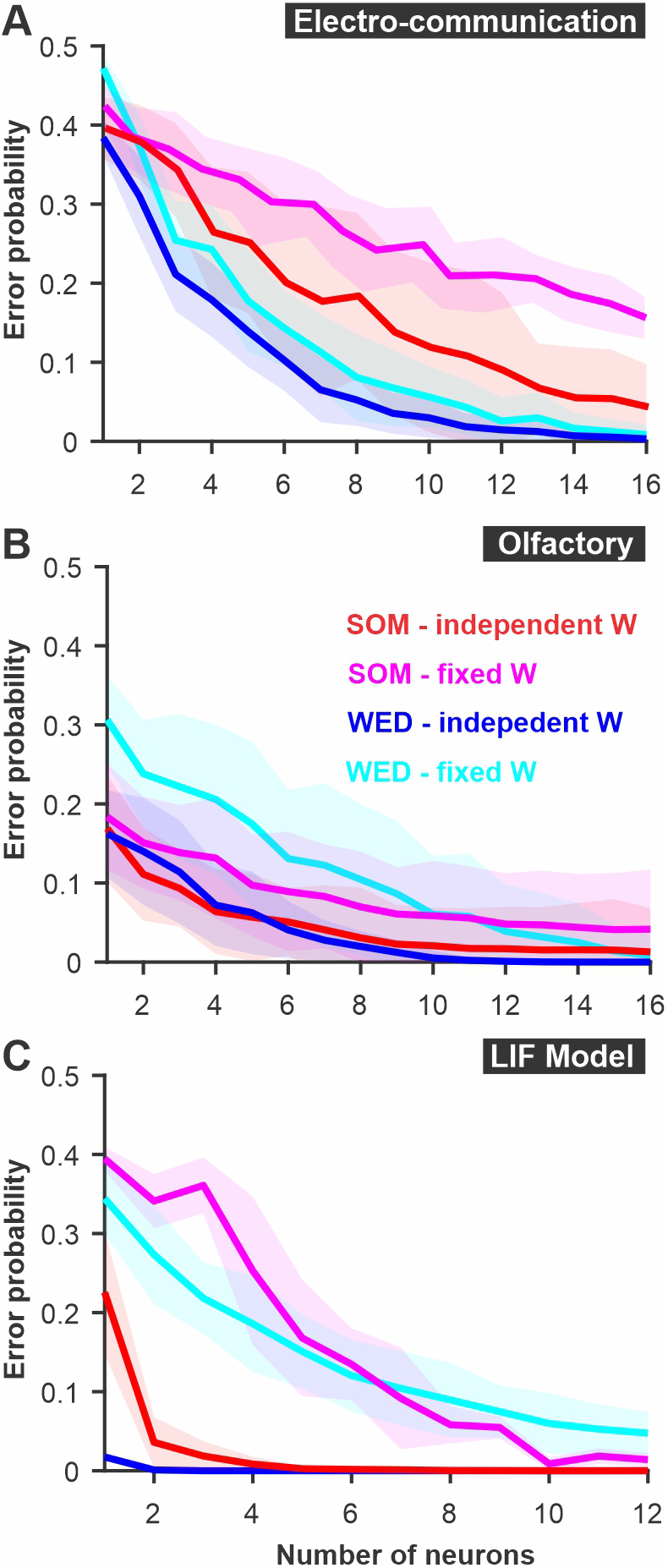
A modified Euclidean distance, where each dimension is weighted, allows accurate discrimination with similar-or better-performance than SOM neural nets. In the “WED” (Weighted Euclidean Distance) analysis, each dimension in Euclidean space is weighted based on the Kullback-Leibler divergence of the response distribution in that dimension. Each dimension (neuron/time bin) can be weighted independently (‘independent W’), or a single weight can be set for a given neuron across time bins (‘fixed W’). Although using independent weights maximizes the information extracted about the difference in stimuli, using a fixed weight emulates a biologically more realistic decoding network. The best method varies across systems: **A.** Electrosensory; **B.** Olfactory; **C.** LIF model. Curves show averages (± s.d.) across all pairs of stimuli.

**Figure 5:**
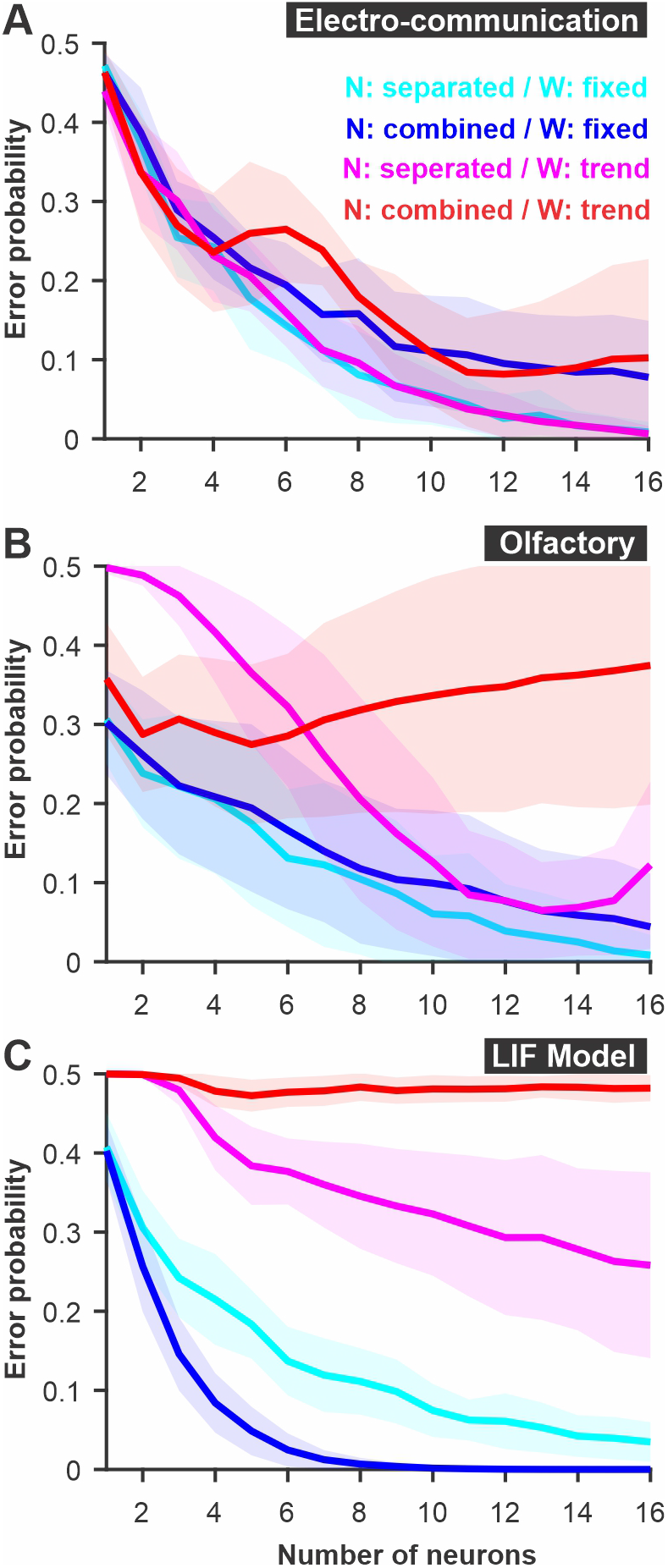
Parameters of a WED analysis can be altered to reproduce biologically realistic constraints on decoding. The different neurons can be kept as separate dimensions in Euclidean space (‘N: separated’) or combined (i.e. average population response) mimicking the fact that synaptic inputs could combine in the postsynaptic terminals of decoding neurons before further processing (‘N: combined’). Weights are calculated independently for each time bin (see Fig 4) and averaged across time to be kept fixed for a given neuron (‘W: fixed’). Alternatively, weights can be fitted to follow a specific function (‘W: trend’). Examples of functions that could be implemented include rules that would replicate firing rate-based plasticity such as facilitation and depression. A firing rate-dependent change in weight across time was not beneficial in the case of the three model systems examined here: **A.** Electrosensory; **B.** Olfactory; **C.** LIF model. Curves show averages (± s.d.) across all pairs of stimuli.

**Figure 6:**
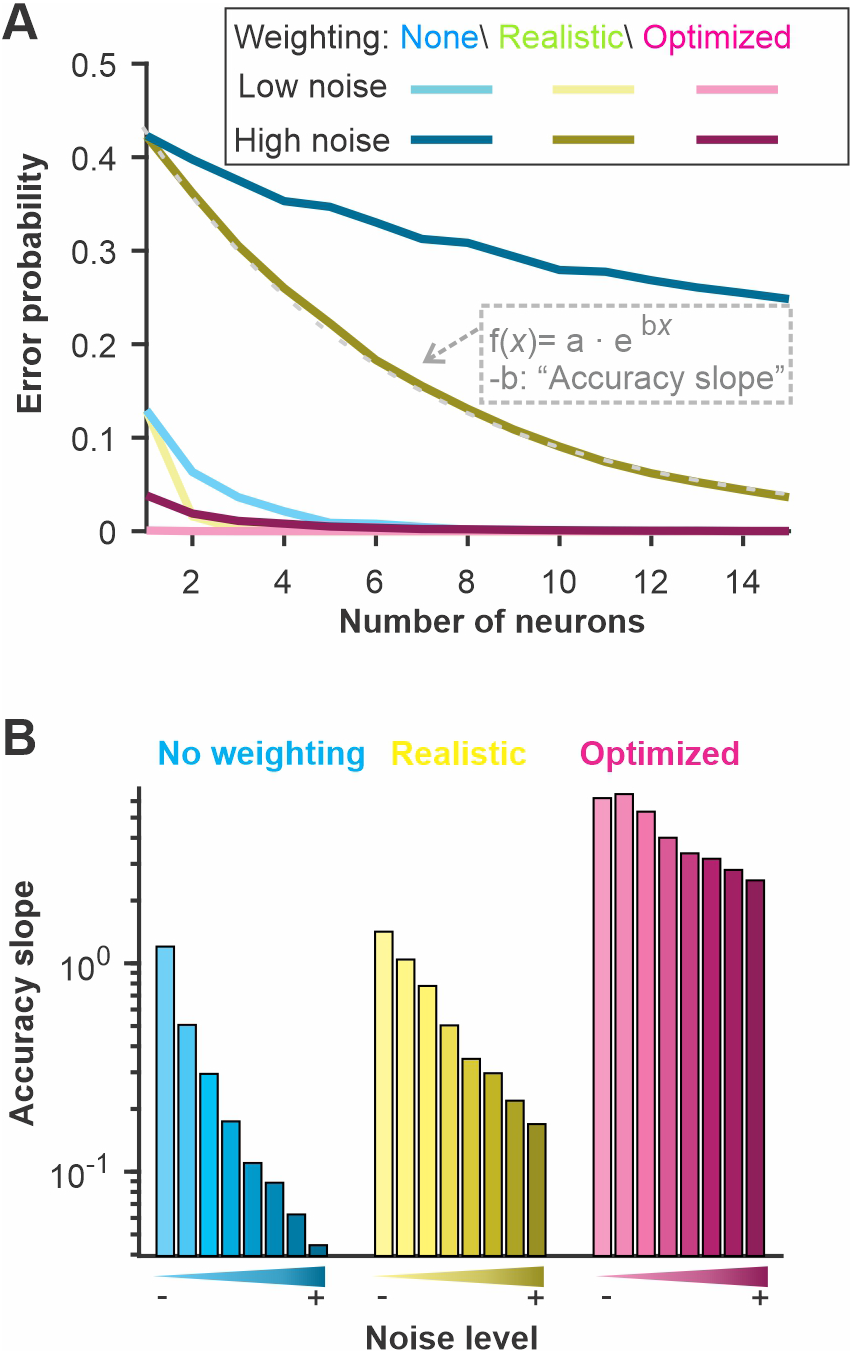
The discrimination accuracy correlates with the signal-to-noise ratio of the neural responses. Varying the amount of noise in our LIF model we quantified the discrimination accuracy using 3 methods. “No weighting”: Euclidean distance with unweighted dimensions and each neuron kept in separate dimensions; “Realistic”: Each neuron is weighted with a fixed weight across time and the different neurons are combined in a single dimension; “Optimized”: Both neuron and time bins are weighted independently and kept as separate dimensions. The “Accuracy slope” is the slope of the exponential fit for the curves displayed in A and therefore reflects how quickly the error level decreases as more neurons are included in the analysis. A sharp decrease (large absolute value of slope *b*) indicates a more accurate coding of stimulus identity. We show: **A**. the discrimination error probability for a large and a small amount of noise as a function of the neural population size used for the analysis and **B.** the derived accuracy for 8 different noise levels.

**Figure 7:**
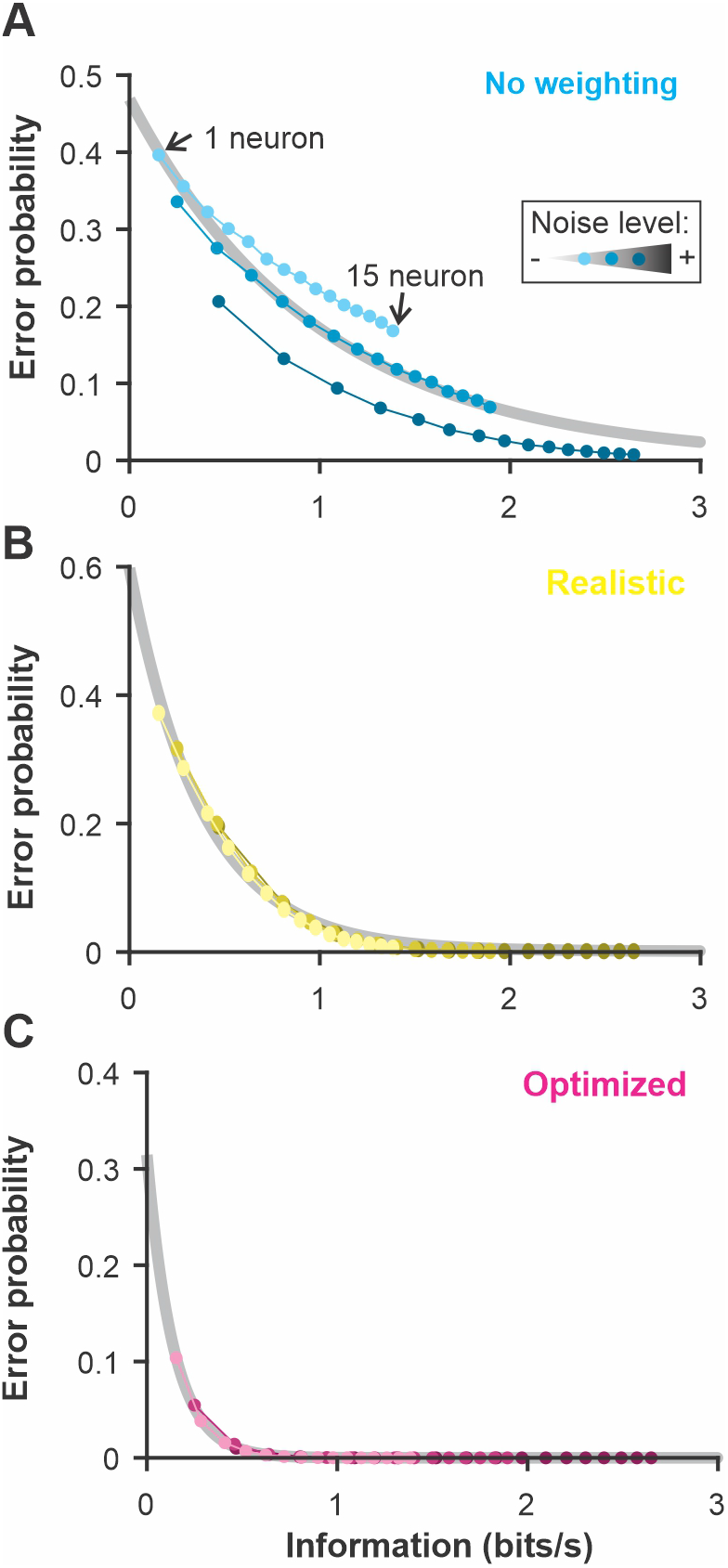
The discrimination error probability is proportional to the information content of the population. The coding accuracy was calculated for this population of LIF neurons using information-theoretic tools. Using the same neurons and stimuli, we calculated the discrimination accuracy (error probability) using the 3 methods described in Fig 6: **A**. Constant weights across neurons and time points; **B.** Weights fixed across time and neural responses averaged across neurons before comparison; **C.** Weight optimized independently across neurons and time points and all the dimensions kept separate. The number of neurons included in the analysis was varied as in previous figures (the different points along each curve here) and the noise in the LIF model was set to 3 different levels (varying shade). The exponential relationship between error and the information content (grey best-fit lines) shows the high correlation between these two measures of coding accuracy (adjusted R-square > 0.98 for B and C).

**Figure 8:**
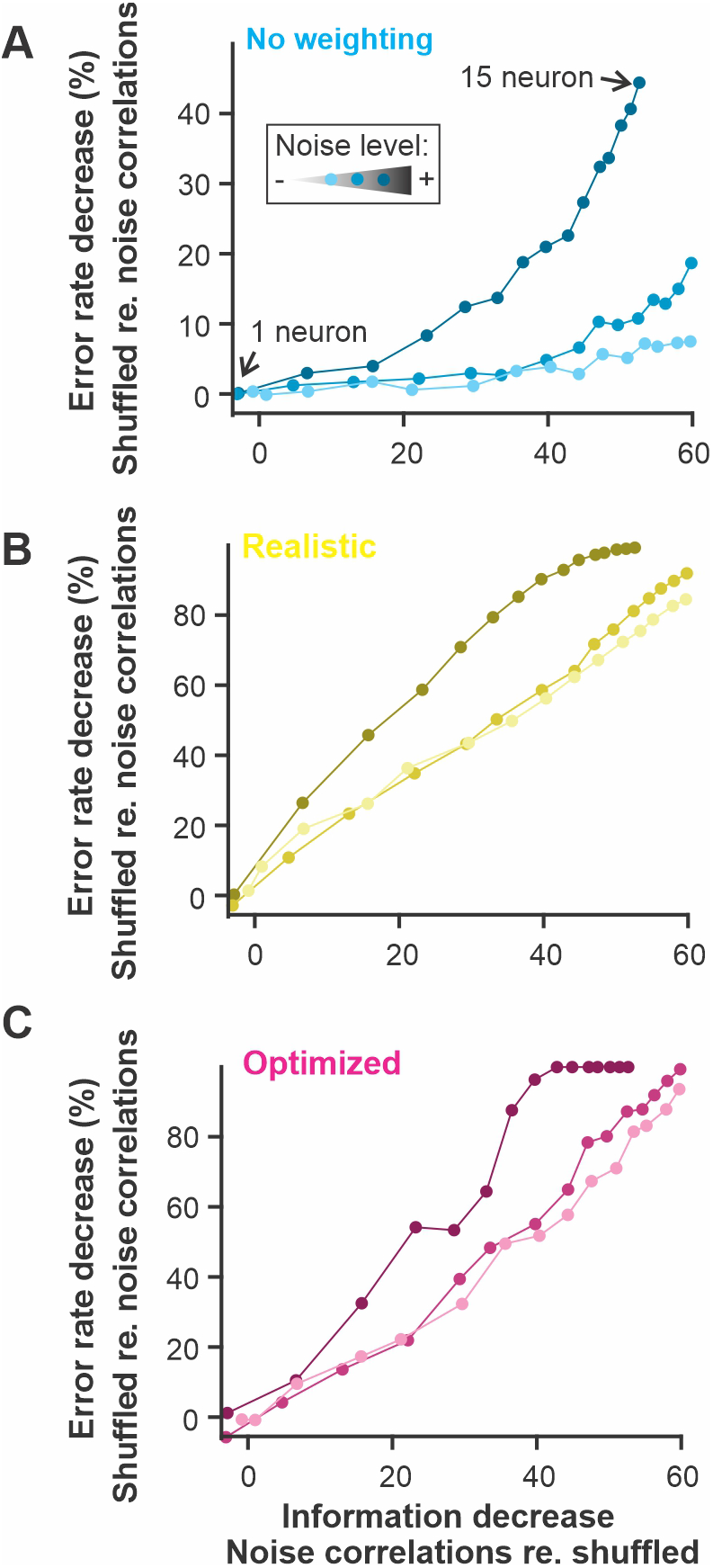
The presence of noise correlations decreases the information content and the discriminability estimate proportionally. Noise-induced correlation across responses of different neurons in a population can decrease the total information content of the population compared to similar responses where the noise is not correlated. Our population of model neurons showed this effect since the population with noise correlations had up to 60% less information that the same responses for which noise correlations were shuffled out. This change in information content is mirrored by a decrease in discrimination accuracy. The figure shows the results for the 3 versions of the analysis used in previous figures: **A.** Constant weights across neurons and time points; **B.** Weights fixed across time and neural responses averaged across neurons before comparison; **C.** Weight optimized independently across neurons and time points and all the dimensions kept separate. The number of neurons included in the analysis was varied as in previous figures (the different points along each curve here) and the noise in the LIF model was set to 3 different levels (varying shade).

### Euclidean distance

Neural responses were expressed as instantaneous firing rate *r* as detailed above. Each population response to a single repeat of the stimulus is a data point in Euclidean space where the number of orthogonal dimensions is *n*·*t* (for a population of *n* neurons and responses with *t* time points). For some analysis (labeled “neurons combined” in the results), the firing rate of the responses of the n neurons was averaged together. The Euclidean distance between pairs of population responses *a* and *b* is:

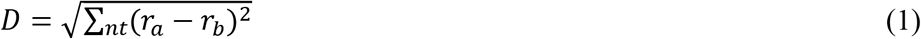

For additional examples of the use of this measure in analyzing sensory responses see Daly et al. (2004a).

### *van Rossum spike distance* (van Rossum, 2001)

Neural responses were expressed as instantaneous firing rate *r* as detailed above and averaged across the *n* neurons included in the analysis to give the population response R for a single repetition of the stimulus. The distance between population responses *a* and *b* is:

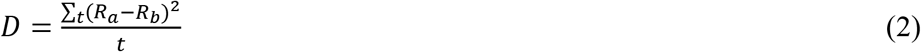

For additional examples of the use of this measure in analyzing sensory responses see Marsat and Maler (2010).

#### Error probability and ROC analyses

To calculate the error probability in discriminating responses to stimulus *B* from responses to stimulus *R* (see red and blue elements in Supplementary Information Figure 2), we build distributions of distances (Euclidean distance or van Rossum distance) for pairs of responses to the same stimulus, *D_RR_* and for pairs of responses to the different stimuli *D_RB_* giving the probability distribution *P*(*D_RR_*) and *P*(*D_RB_*). Since we have responses to many repeats of the stimuli from many neurons there are many possible combinations of responses (particularly if the responses of the different neurons are not recorded simultaneously), Therefore we take a random subset of combinations. For example, consider that we recorded separately the responses of 20 neurons to two different stimuli and presented each stimulus 30 times and we want to quantify the discriminability based on a subset of 3 neurons. To build the probability distribution *P*(*D_RR_*) and *P*(*D_RB_*), we could take 100 different combinations of 3 randomly selected neurons out of the 20, and for each combination, we take 100 combinations of repeats giving 10,000 different pairs of population responses to build the probability distribution. If the different neurons are recorded at the same time (thereby possibly containing noise correlations), the same repeat must be taken across the neurons of a given combination.

Receiver operating characteristic curves were generated by varying a threshold distance level T. For each threshold value, the probability of detection (P_D_) is calculated as the sum of *P*(*D_RB_*>*T*), and the probability of false alarm (P_F_) as the sum of *P*(*D_RR_*>*T*). The error level for each threshold value is E = 1/2P_F_ + 1/2(1 - P_D_). The discrimination error levels reported in the figures are the minimum values of E.

Alternatively, the distribution P(*D_RB_*) can be based on the distance calculation between a given stimulus *R* and all the other stimuli in the data set. This approach is more similar to a confusion matrix analysis (for an example of confusion matrix used in a similar analysis, see Chase and Young, 2006). Error rates will then depend on the ensemble of stimuli present rather than on pair-wise comparisons. In many cases, we find it more useful to obtain an accuracy number for a single pair of stimuli and so we only present this approach here. We also note that the ROC analysis could be based on having an upper and lower threshold T rather than a single one; this strategy was not explored. This method results in an estimate of error probability that is based on the distributions of responses and therefore has the characteristics of a statistical analysis so no further statistical analyses were performed unless otherwise noted.

#### ANN: SOM neural net

In choosing a type of ANN to use for decoding neural responses a variety of strategies are available: recurrent network vs. non-recurrent, deep vs. shallow; supervised vs. unsupervised, etc. Our goal is not to find the most performant type of network for the task but on the contrary to determine how the simplest type of network would perform. In doing so we will be able to argue that the most basic feature of ANNs -weighing of inputs-provides a powerful approach to decoding neural responses. We, therefore, opted for a shallow, unsupervised, and non-recurrent type of neural network the Self Organizing Map (Kohonen, 1982).

We used the SOM neural net tool built into Matlab 2017b (Mathworks, Natick, MA) and detailed documentation can be obtained from the provider. Note that it can be accessed through the Neural Network Clustering app in more recent versions of Matlab (e.g. 2022b). Briefly, the network has a single competitive layer of *X* inputs (here as many inputs as neurons x timepoints) connected to a small number of *Y* output nodes (here we use 2 output nodes for our pair-wise stimuli discrimination). During the training phase, each set of inputs (a population response) is attributed to the output node for which the connection weight has the smallest Euclidean distance with the input pattern. The weights of the output node are then shifted to be closer to the datum it received. As the network is trained with a subset of data, it thus learns to classify the data in *Y* clusters by maximizing the Euclidean distance between the clusters. A different subset of data is used for testing. We quantify the performance of the classification by attributing each output node to a stimulus (the stimulus for the majority of the responses attributed to the node) and calculating the proportion of miss-classified inputs (with 0.5 representing chance-level because each node has received an equal number of inputs from both stimuli). This process is illustrated schematically in Supplementary Information Figure 1.

#### WED method

Our Weighted Euclidean Distance analysis consists of the same sequence as described above: preparation of the population responses by convolving each spike train with an alpha or gaussian function, Euclidean distance calculation, and ROC analysis. In addition, we weigh each dimension of the Euclidean space by multiplying the firing rate in that dimension with a weight *W_nt_* before the Euclidean distance calculation. Weights are calculated as follows.

The method used in all main figures of this paper calculates an optimal weight associated with each dimension based on the Kullback-Leibler (KL) divergence (Kullback and Leibler, 1951). It is a measure that quantifies the information required to characterize the second distribution if we know the first one; it was described by Kullback as a “discrimination information” (Kullback et al., 1987). The probability distributions *P_Ri_* and *P_Bi_* of the firing rates *x* in responses to stimuli *R* and *B* in each of the *i* dimensions (*number of dimensions* =*n*·*t* for a population of *n* neurons and responses with *t* time points). The KL divergence is calculated as:

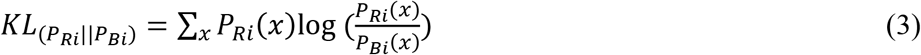

Note that the KL divergence for 2 distributions is not symmetrical: *KL*_(*P_Ri_*||*P_Bi_*)_ ≠ *KL*_(*P_Bi_*||*P_Ri_*)_. However, when we perform the ROC analysis, we use the responses to one stimulus to build the intra-stimulus distance (e.g. *P*(*D_RR_*)) to compare to the inter-stimuli distance (*P*(*D_RB_*)) we thus take the corresponding KL divergence (e.g. *KL*_(*P_Ri_*||*P_Bi_*)_) and repeat the analysis with the reciprocal comparison.

KL values are normalized, by dividing by the average KL value, to obtain weights with an average value of 1 for each distance calculation:

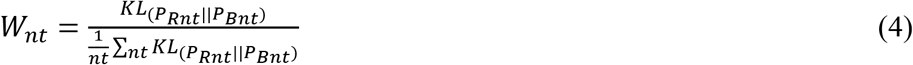

In another version of this analysis (see Supplementary Information Figure 3) the weight is based on the area difference *AD*(*_P_Ri__*,*_P_Bi__*) between the two distributions:

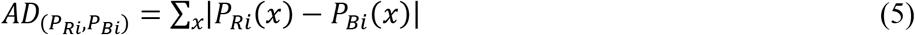

AD values are normalized to 1 by addition:

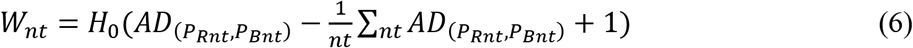

where H_0_ is the Heaviside step function. Results using this method of calculating weights are similar to results using the method based on KL divergence so only the latter are presented in the main text.

#### Information measure

Information content of LIF responses was evaluated using well-established methods of information estimates based on coherence measures (Borst and Theunissen, 1999); see also (Marsat and Pollack, 2004; Marsat et al., 2009) for additional details. To do so, responses from different neurons were averaged and stimulus-response coherence was calculated. Upper-bound coherence was also calculated using the response-response coherence but since our responses are linear, there was no difference between upper- and lower-bound coherences (see Supplementary Information Figure 6).

All analysis was performed in Matlab 2017b, 2019a, or 2022b (Mathworks, Natick, MA). The code for the analysis described in this paper and example of source data are available at https://www.marsat.org/publications/code

## RESULTS

Using datasets from different model systems allows us to make more general statements about the performance of the analysis and also to determine how different coding aspects are revealed by the methods. For example, the PN olfactory responses show a clear diversity in response patterns across neurons (Fig. 1) and it is known that the identity of the neurons active for a given stimulus is key in encoding the stimulus (Daly et al., 2016). In contrast, the PC electrosensory responses show qualitatively more similar response patterns across neurons and it might not be critical to evaluate the neuron-specific response pattern to extract all the information. Another example can be given by comparing the temporal pattern of responses between the olfactory and model responses. For model responses, the detailed temporal structure is key in encoding the stimulus identity and the average firing rate is similar across stimuli. For olfactory neurons it is the contrary, the overall firing rate varies largely from neuron to neuron across stimuli but firing rates varies less rapidly across time. The methods discussed in this paper can be used to demonstrate and quantify differences in coding strategies such as these.

### Established spike-distances methods

We first compare two established ways to quantify differences between spike trains: the Euclidean distance and the van Rossum spike distance metric (see methods). For both we use a ROC analysis to compare the results and display the discrimination error levels as a function of the number of neurons used for the analysis (Fig. 2). The calculation was performed for each stimuli pair, then averaged; standard deviations displayed shows the variability in the averages across stimulus pairs. We note that, rather than pair-wise comparisons, the analysis can compare the responses to one stimulus to the responses of all other stimuli in a way that would be closer to a confusion matrix (see Methods for additional details) but we do not present results for this method here.

There is an appreciable difference in discrimination performance for the different datasets. The electro-communication stimuli were weak and thus the responses provided a fairly noisy representation of stimulus identity. As a result, the discrimination error levels were high even when the analysis based itself on a population of 16 neurons (Fig. 2A). Olfactory and model responses were less noisy and led to more accurate discrimination (Fig. 2B, C) for analysis based on Euclidean or van Rossum spike distance metrics (red and black curves). For the electrosensory and the olfactory responses, the method based on the Euclidean distance allowed a slightly better discrimination accuracy whereas, for the model responses, the method based on the van Rossum spike distance metric gave significantly better results. The only major difference between the methods, as we implemented them, is that each neuron was kept as a separate dimension in the Euclidean distance whereas responses were averaged across neurons before performing the van Rossum analysis. If neuron identity is important in encoding stimulus identity, keeping the neurons as separate dimensions will allow extracting more accurate information whereas if each neuron is similar in its response pattern, averaging across neurons averages out the noise. To confirm that this is indeed the main difference, we repeated the Euclidean distance analysis after averaging together the responses of the different neurons. We show that discrimination performance is similar to the van Rossum method (Fig. 2) demonstrating that this comparison allows us to assess the importance of differences between neurons in encoding the information at the population level.

### Comparison with an artificial neural net

For both the olfactory and electrosensory datasets, we know that the animal is able to discriminate the stimuli as demonstrated by behavioral assays (Daly et al., 2008; Allen and Marsat, 2018). Even though these systems can rely on neural populations larger than tested here, the poor discrimination performance suggests that we might not be extracting as much information as the system actually does. This seems to be particularly obvious for the responses to electrocommunication stimuli that remain at chance levels for all population sizes tested. We, therefore, tested a method that has proved to be very efficient in clustering patterns: artificial neural networks. Specifically, with used a SOM-type network that relies on unsupervised learning to change the way it weights inputs to optimally cluster the data into a fixed number of outputs (see methods).

The SOM network performed much better than the other two methods described above (Fig 3). As a result, discrimination was nearly errorless in all three systems for the larger population sizes. This result suggests that the core principle used by SOM networks -differential weighting of inputs-allows much more efficient decoding of the responses. This result is not surprising and can be understood by the fact that neurons or time points that are very noisy and contain very little information about stimulus identity will be weighted down and thus will not influence (i.e. add noise) the discrimination process. More complex neural networks with more layers (i.e. deep neural nets) and a supervised learning component could potentially perform even more reliably in these discrimination tasks. However, the convenience of such analysis methods also decreases as the network complexity increases. Several factors limit the convenience of neural networks in analyzing sensory responses. First, the time required for running the analysis on regular computers becomes unreasonably long when responses with many dimensions (time point X neurons) are used. To illustrate this point, using a computer with 10 cores, the SOM analysis in Fig. 3C took several days to run. Also, for more complex networks, larger datasets might be required. Moreover, deep neural networks perform a transformation on the input that is more opaque to the experimenter and that would require tedious secondary analysis to detail once the network has learned its proper configuration. To benefit from the advantages of weighting the inputs exploited by the neural nets but keeping the analysis simple we enhance the spike distance metric with a weighting procedure.

### WED measure

In our Weighted Euclidean Distance (WED), we aim to use the Euclidean distance but weigh each dimension according to how informative it is about the difference in the stimulus. In other words, the weight has to be proportional to how distinct the distributions of responses are in that dimension. We tested two methods to weigh inputs. The results displayed here are based on the Kullback-Leibler divergence (Kullback and Leibler, 1951) and the weights are then normalized by dividing with the average weight across dimensions. This normalization is essential to keep the overall weight of the inputs unchanged. We also tested a weighting procedure based on the integral of the difference between the two distributions normalized to one by an additive procedure (see methods). This second method is qualitatively similar to the one presented here (see Supplementary information, Fig 3).

The WED method performs as well or better than the SOM neural net in permitting discrimination of the stimuli (red and blue curves, Fig. 4). We note that both analyses rely on weights that can change abruptly from one time point to the next in ways that are not dictated by biologically realistic rules. Indeed, the analysis is aimed at quantifying how well an ideal observer, with sufficient prior knowledge, could discriminate between two stimuli based on the pattern of neural responses. It does not explicitly emulate how well a biologically realistic decoding network could perform. We can, however, adjust the way the weights change as a function of time to replicate more biologically plausible mechanisms. Various time-dependent or firing-rate-dependent mechanisms replicating rules of synaptic plasticity could be implemented. In figure 4 we explore the simplest rule: keeping synaptic weight fixed across time. We average the KL values across time for a given neuron, thereby each neuron is simply weighted according to its average information about the difference between two stimuli. We can see that the difference between independent and fixed weights across time varies for the 3 model systems tested here (Fig 4). This indicates that in some responses (e.g. model), information about stimuli differences varies across time whereas for others (e.g. olfactory) the informative dimensions are fairly constant in time.

In figure 5, we present two additional alterations to the WED methods that replicate biologically-inspired constraints and that can clarify how an efficient decoding network could be designed and what aspects of the responses carry useful information. We implement a short-term plasticity rule that allows the weights to change as a function of the preceding firing rate mimicking short-term facilitation and depression. The rule is adjusted to fit the changes in weights determined independently at each time point within the constraint of biologically plausible dynamics (see Methods for more details). Implementing such a rule could lead to improvement of discrimination performance in the analysis if the biological system is built to encode and decode through such mechanisms. In our case the procedure did not lead to improved discrimination (compare cyan and magenta curves, Fig 5), suggesting that the rule we implemented would not offer a decoding benefit over having fixed weights over time. We also revisited a concept introduced at the beginning of the results section: having the different neural responses combined or kept as separate dimensions. A postsynaptic decoder could receive inputs from all neurons in the population and the postsynaptic potential be combined in the decoding neurons before any further processing occurs. Averaging responses across neurons before population responses are compared would replicate this scenario (blue curves, Fig 5). Alternatively, a specific subset of neurons connects to a decoder, and the identity of which neuron is active at what time determines the activation of the decoder. To replicate this decoding scenario, we keep neurons as separate dimensions in the analysis (cyan curves, Fig 5). Our results show that some systems would benefit from the former procedure (Fig 5C) and others from the latter (Fig 5 A&B).

We note that an informative aspect of these types of analysis (both our WED decoding measure and the SOM-based measure) is that weight patterns can be analyzed. For example, we visualized the strength of the weights for the odor discrimination analysis and we noticed that for each discrimination task 7-9 different neurons were strongly weighted out of the 16 in our dataset (Supplementary Information, Fig 4). Interestingly, its is believed that the Kenyon cells, which decode the information from PNs and are involved in odor identification, receive inputs from a qualitatively similar number of PNs (Jortner et al., 2007; Honegger et al., 2011). The input to the decoder can be manipulated in various other ways, for example by including a longer or shorting portion of the recording after the start of the stimulus (Supplementary Information Fig 5) thereby assessing how performance would improve as the animal senses the stimulus for a longer period.

### Noise and information

To relate our WED measure to an information-theoretic quantification of coding performance, we ask how noise affects the discrimination accuracy estimated by our method and how it related to the amount of information carried by the spike trains. To do so, we focus on the LIF model responses to be able to change systematically the amount of noise in the system. We also performed the analysis by keeping the noise fixed but changing the signal strength, thus similarly affecting the signal-to-noise ratio, and the results were qualitatively similar (Supplementary Information, Figure 6). We first confirmed that increasing the amount of noise in the model decreased the discrimination performance assessed by our analysis (Fig 6). We use three versions of the analysis: a “basic” analysis with no weighting (i.e. based on a traditional Euclidean distance; cyan curves), a “realistic” analysis where weights are kept constant across time and the responses from different neurons are averaged before the distances are calculated; and one analysis labeled as “optimized” where weights are optimized independently across neurons and time points and all dimensions are kept separate.

The use of white noise stimuli conveniently allowed us to calculate the information coding rate about the temporal modulations by calculating the coherence between stimulus and response or the response-to-response coherence (Borst and Theunissen, 1999). Since we used a linear system in this analysis, the two types of coherences lead to similar estimates of information rate (Supplementary Information, Figure 6). Information rate was calculated for population responses of varying sizes (similar to the analysis in previous figures) and neurons with different amounts of noise. By plotting the discrimination accuracy as a function of the information rate calculated for the same responses (Fig. 7), we show that the 2 are highly correlated. These results demonstrate that the discrimination performance of these analyses is directly related to the information content of the spike trains but that the analytic procedure dictates how efficiently the information is used to discriminate between two stimuli.

### Noise correlations and population responses

Correlations among neurons of a population can be an important aspect of the response affecting the information it carries about the stimulus (Kohn et al., 2016). Stimulus-elicited correlations will obviously be taken into account by the WED method (or other spike distance metrics) since correlated vs uncorrelated responses will lie in different clusters of multidimensional response space. It is less obvious that noise correlations will influence the result of the analysis. When common sources of noise cause variability in neural responses that are not related to the stimulus, this correlated noise decreases coding performance -as would any type of noise-but it cannot be averaged out across these neurons since they contain the same noise. In most cases, a population with correlated noise in their responses will therefore carry less information than a population with similar but uncorrelated noise (Kohn et al., 2016; Hazon et al., 2022). To address this issue, we introduced noise correlations in the population of model responses. As expected, these noise correlations decrease the information content of the population compared to responses without noise correlations. More importantly, a proportional decrease in discrimination accuracy was observed (Fig 8). This comparison (shuffled vs un-shuffled) can therefore be applied to neural responses recorded at the same time to gauge if correlated noise affects discrimination performance. Further, these results demonstrate that the analysis can be applied to a wide range of neural response types even if they include noise correlations that significantly affect their encoding accuracy.

## DISCUSSION

We compared different methods of quantifying the accuracy with which a population of neurons encodes the identity of the stimulus. We show that the estimate of the discrimination performance of a system varies widely based on the method used. Methods that weight inputs provide a clear advantage since they can base the results on information-rich portions of the population response while de-emphasizing noisy portions. A concrete illustration of the principle would be the visual discrimination of twins. Differences in their facial features might be so subtle that they are within the noise level of our perceptual judgment. However, specific features (haircut, a freckle…) might allow reliable discrimination and a classifier should consider this feature above all else. Decoding networks in actual biological systems can rely on the same principle, weighting more heavily the input from specific neurons (e.g. specific sub-populations) or certain time frames (e.g. the beginning of a call) of the population response.

We also argue that our WED decoding method can be adjusted in several ways to examine how sensory information is encoded in the population response. Inputs from different neurons can be combined or kept separate to understand whether the encoding of stimulus identity is distributed across space (different combinations of neurons being active for different stimuli). We showed that the effect of noise correlations on population coding could be assessed by comparing population coding in responses that were recorded at the same time vs shuffled responses. We can also identify the subset of neurons that a decoder would need to be connected with to achieve accurate discrimination by analyzing the weight pattern of the decoder.

The contribution of temporal resolution to coding can be assessed by combining time bins or smoothing the spike trains (not shown in this paper but see Marsat and Maler, 2010). The discrimination performance of the decoder can also be compared to the latency of the behavioral response. By including longer or shorter windows of the recording after the start of the stimulus, we can evaluate how the putative discrimination performance would improve as the animal processes the stimulus for a longer period. The relevance of spike patterns can also be investigated by allowing weights to change over time or not. Particularly, specific rules can be implemented to change the weight across different time points in a way that emulates biologically realistic processes. For example, various firing-rate-dependent weighting rules can be considered can be implemented (e.g. see Lappalainen et al., 2019). By plotting the independently-determined weight against the average firing rate in the preceding time points, we can determine the best-fit function relating the two and implement the function to determine the weight at each time point. Although our dataset did not benefit from such a decoding procedure, we suggest that this method can help identify instances where this decoding procedure would be beneficial. It should also be noted that, although we limited this paper to using firing rate in narrow time bins as the input, the same method can be implemented using derived measures of the neural response such as phase-locked spike rates, coincident spike per seconds, gain, or any other measure of neural respond that could be relevant to sensory coding.

Of the various methods available, including the ones we considered in this paper, some provide theoretical measures of information content (e.g. coherence based information theoretic analysis as in: Chacron et al., 2003) while others excel in accurately decoding the stimulus identity (e.g. deep neural network decoding as in: Wen et al., 2018)). We are particularly interested in decoding methods that give us insight into the biological process involved in decoding neural responses. The analysis we demonstrate in this paper -the WED decoding method-starts with convolving the spike trains with an EPSP-like filter mimicking the process a post-synaptic decoder would implement. Response patterns are then compared in a way that could be realistically implemented by neurons (Larson et al., 2009). The way the responses are weighted and combined can be adjusted to replicate various synaptic integration principles (see previous paragraphs). Lastly, the method outputs a measure that is easy to relate to behavioral performance: the probability to make an error in discrimination. We argue that the method is intuitive to use for neuroscientists, flexible, efficiently extracts the information encoded in the spike trains, and does so in a way that is biologically realistic.

## Acknowledgments

Funding was provided by the National Science Foundation grant IOS-1942960. We acknowledge E.M. Staudacher for his role in collecting the published dataset of olfactory responses in moth collected while working in the Daly laboratory – a dataset that is used in this paper for testing our analyses.

## Conflict of interest statement

The authors declare no competing financial interests.

## Supplementary Information

**Supplementary Information Figure 1:**
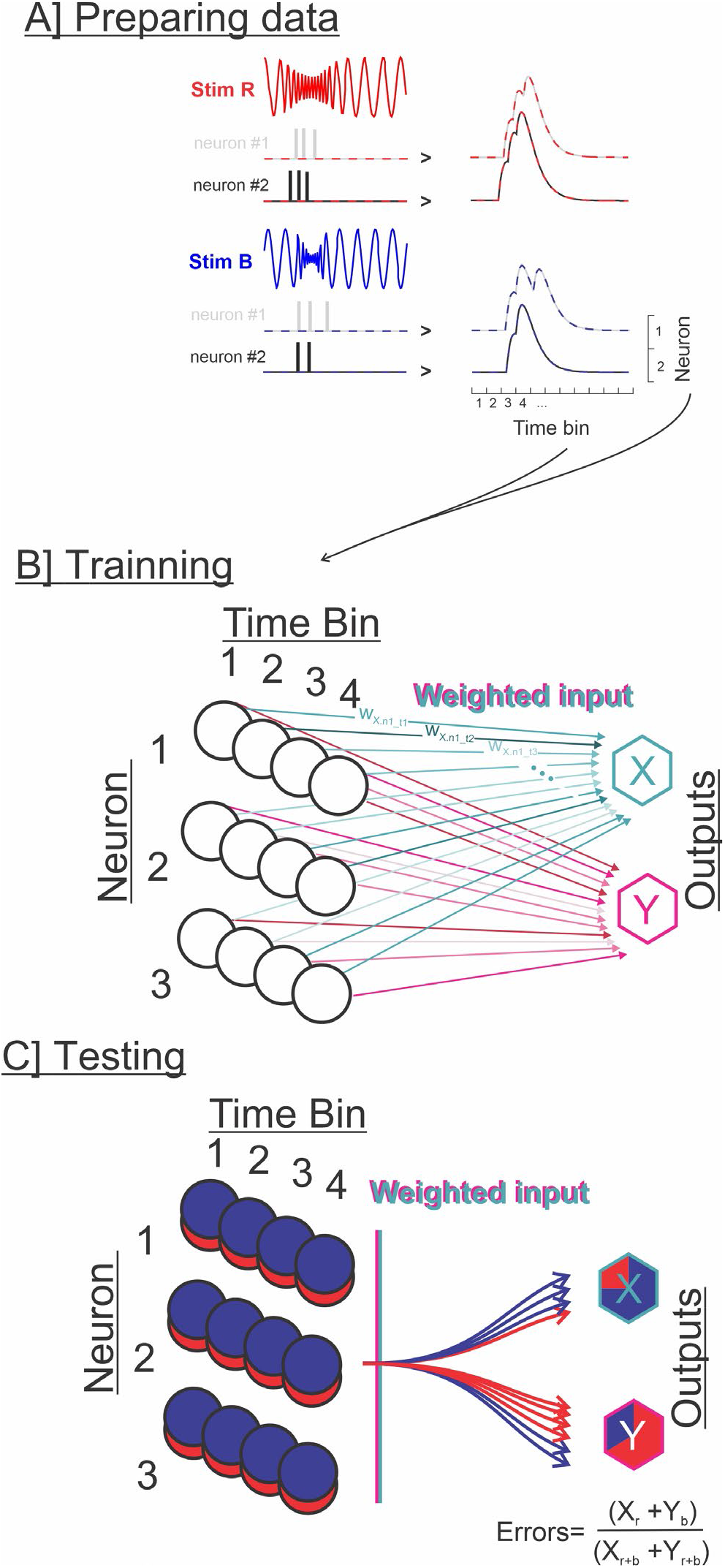
Using a SOM neural network to test responses discriminability. **A.** Spike train responses to two different stimuli are prepared the same way it is in the other analysis used in this paper: binary spike trains are convolved with an alpha function (resembling the time course of post-synaptic potentials) resulting in an estimate of instantaneous firing frequency. The firing rate for each neuron and at each time point can then be used as inputs to the SOM neural netThe Self Organizing Map (SOM) neural network available in Matlab 2017b (Mathworks, Natick, MA) is a shallow unsupervised neural network with an input layer and an output layer. The weights between the input and output nodes are optimized based on the variability of the input patterns to maximize the clustering in separate groups. We use networks with the same number of inputs as dimensions in our population responses (the product of [number of time points] x [number of neurons]). The number of outputs is equal to the number of stimuli; here we tested pair-wise discrimination so we have 2 output nodes. **C.** After training, a different subset of data is used for testing. Classification accuracy is calculated, where an error probability of 0 is reached if all the responses to the two stimuli as classified in separate outputs and 0.5 error probability indicating that each output receives data point from either stimulus equally (chance level).

**Supplementary Information Figure 2:**
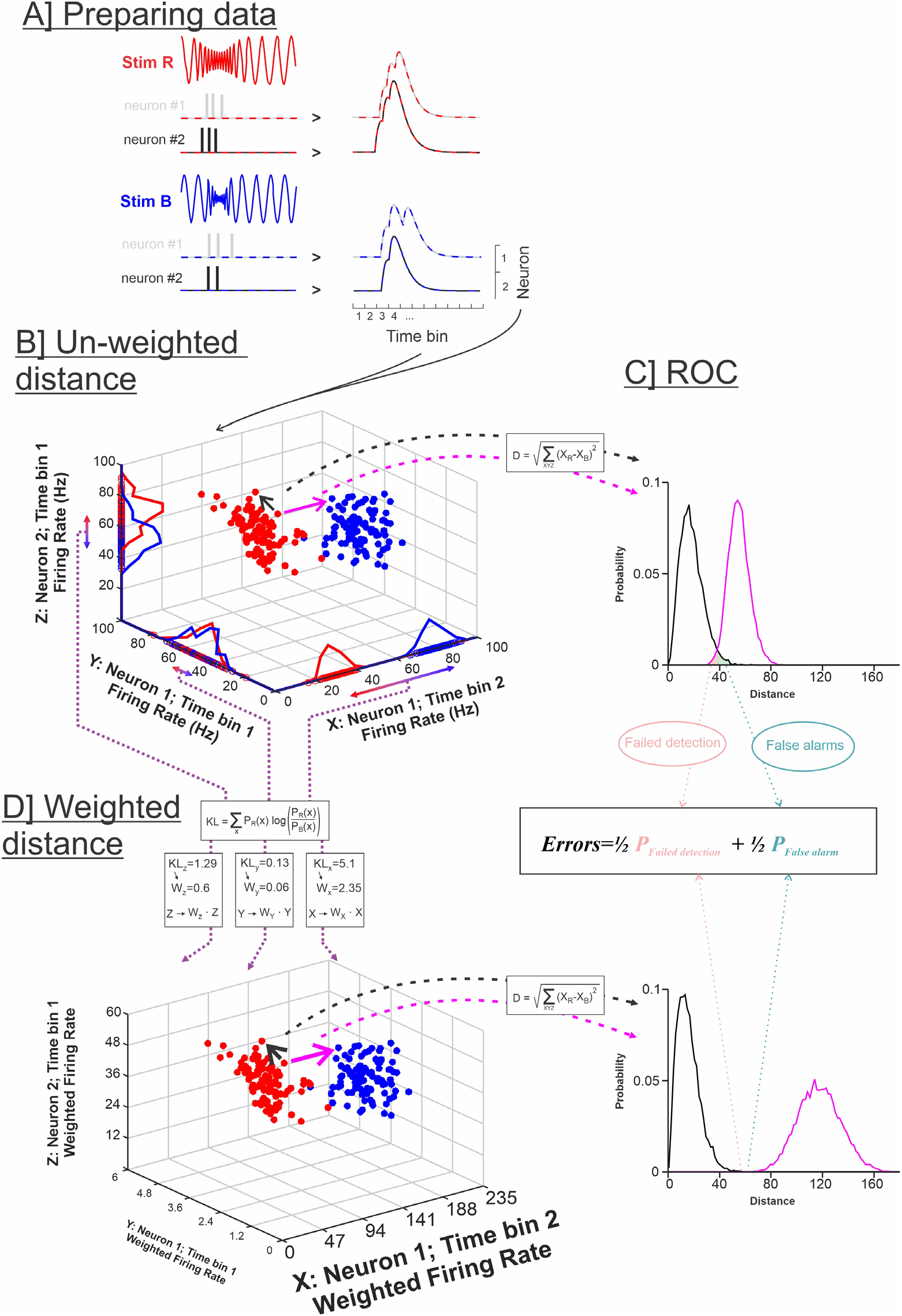
Weighted and unweighted Euclidean distance analysis. **A.** Spike train responses to two different stimuli are prepared as for other analyses and replicates the procedure used in van Rossum (2001). Binary spike trains are convolved with an alpha function (resembling the time course of post-synaptic potentials) resulting in an estimate of instantaneous firing frequency. The firing rate for each neuron and at each time point can then be used as inputs to calculate the spike-train metric distance. Each population response is taken as a data point in multi-dimensional Euclidian space with each orthogonal dimension representing the firing rate of one neuron at one time point. Therefore the number of dimensions is the product of [number of time points] x [number of neurons]. We depict 3 dimensions here for illustration purposes. A large random subset of pairs of responses (to the two stimuli – red and blue) is compared by calculating their Euclidean distance (magenta). Considering the number of stimuli pairs and the number of repeats, there are often millions of possible pair-wise comparisons, therefore, requiring a random sub-sample of combinations to be taken. Pairs of responses to the same stimulus are also compared (black Euclidean distance) so we can compare the inter-stimuli to the intra-stimulus distances. The inter- and intra-stimulus distances are expressed as distributions (magenta and black distributions respectively). The overlap between the distributions can be quantified, assuming an ideal observer, using a ROC analysis to obtain a minimum error probability. D. When proceeding to weigh each dimension in the Euclidean distance analysis, the overlap in the distribution in each dimension is quantified. The distributions in each of the 3 dimensions displayed in B are displayed along the analysis and we can see that some overlap almost completely (e.g. in dimension Y) whereas some do not overlap at all, (in dimension X) leading to small and large Kullback-Leibler divergences respectively (see equation and numbers in the boxes). The weights are the result of a normalization (dividing by the average KL divergence across all dimensions) of the KL values so that the average weight is 1 for each pair-wise comparison. Euclidean distance is calculated after multiplying the firing rate in each dimension by the corresponding weight (notice the change in axis scale between the 3D plots in B and D). The overlap between the distributions for intra- and inter-stimuli Euclidean distances is thereby reduced (see bottom-right plot used for ROC analysis) and the discrimination error probability is minimized.

**Supplementary Information Figure 3:**
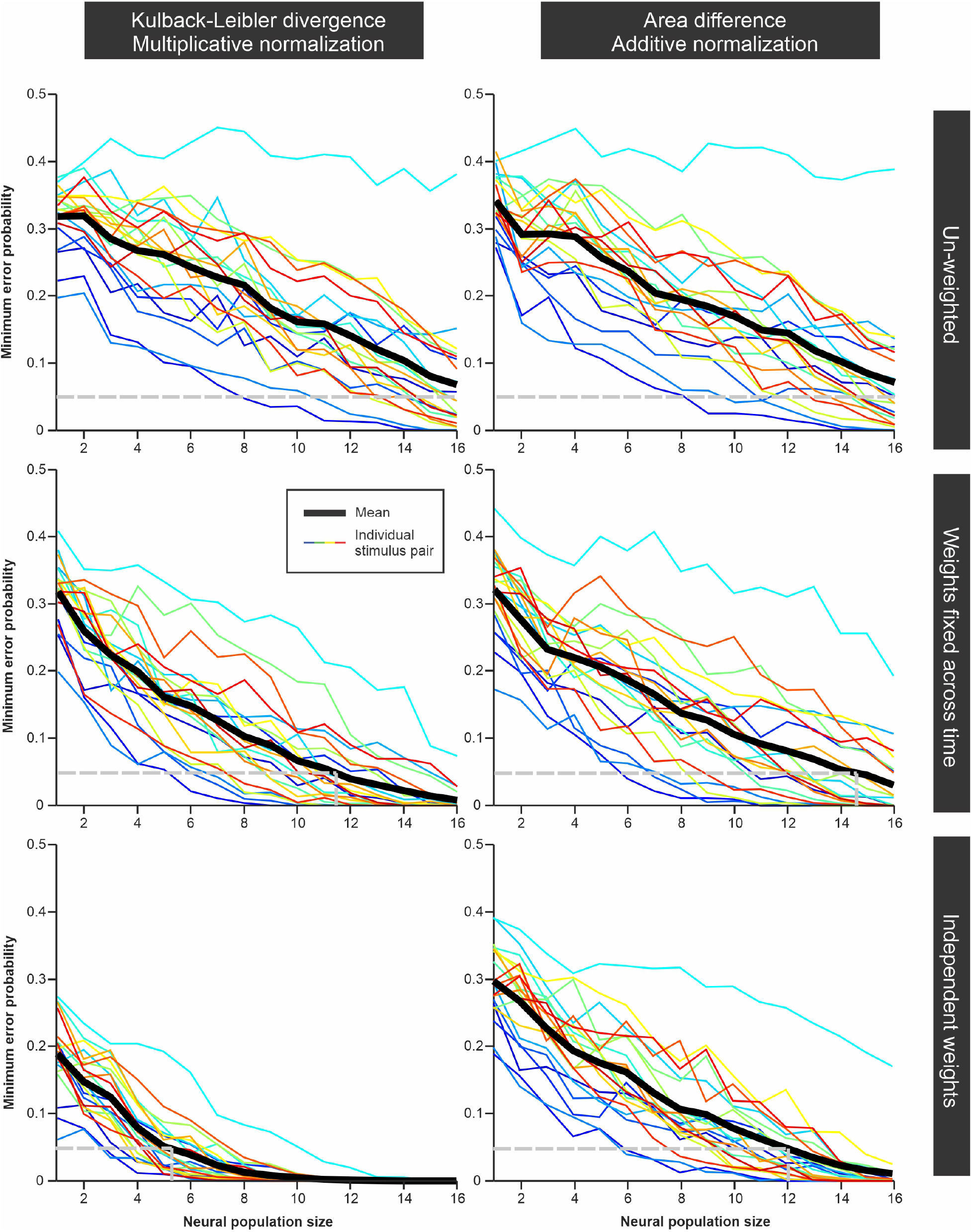
Comparison of the WED discrimination analysis using two different weighting methods. In the plots in the **left column**, we calculate the Kullback Liebler divergence for the distributions of firing rates in each dimension for a given stimulus pair (see Supp Fig 2). These KL-based distances are then normalized by dividing with the mean KL-based distances across dimensions thereby giving an average weight of 1 in each comparison. In the **right column**, we quantify the overlap between distributions by taking the absolute value of the differences between the two distribution curves (this value will vary between 0 and 2 since each distribution has an integral of 1). The mean -across dimensions-of these overlap measurements is then shifted to 1 (by adding 1-mean) and any negative values are set to 0. The two weighting methods are fully implemented in the two **bottom plots**. In the **middle plots,** the weights calculated independently for each dimension are averaged across time points for each neuron in the population before calculating the Euclidean distance (i.e. one neuron=one weight). In the top plots the different dimensions are not weighted (all weights =1) thus the two plots differ only slightly reflecting the difference in the randomly selected pairs of stimuli/repeats used in the analysis. The plots displayed here are based on the olfactory responses to 7 odors and each of the 21 pair-wise comparisons is shown in a different color with the average across stimuli pairs in black. We highlight with a grey dashed line the 0.05 error probability, and the average number of neurons required to reach this reliability level, as a simple way to compare discrimination accuracy across datasets/analysis. It is interesting to note that the KL-based analysis outperforms the area-based method, particularly when weight can vary in time (bottom two plots). We have observed, however, that for some datasets (data not shown), the more pronounced weighting of our KL-based method led to results that were maybe unrealistically good. Due to the multiplicative normalization, this method can attribute very strong weights to just a few time points/neurons with the other weights being much weaker. It is unclear whether biological systems could implement such extreme weighting, particularly across time. More biologically realistic estimates could therefore be based on the method presented on the right (where weights are normalized by an additive process and thus vary less strongly across dimensions) and/or by keeping weights constant across time.

**Supplementary Information Figure 4:**
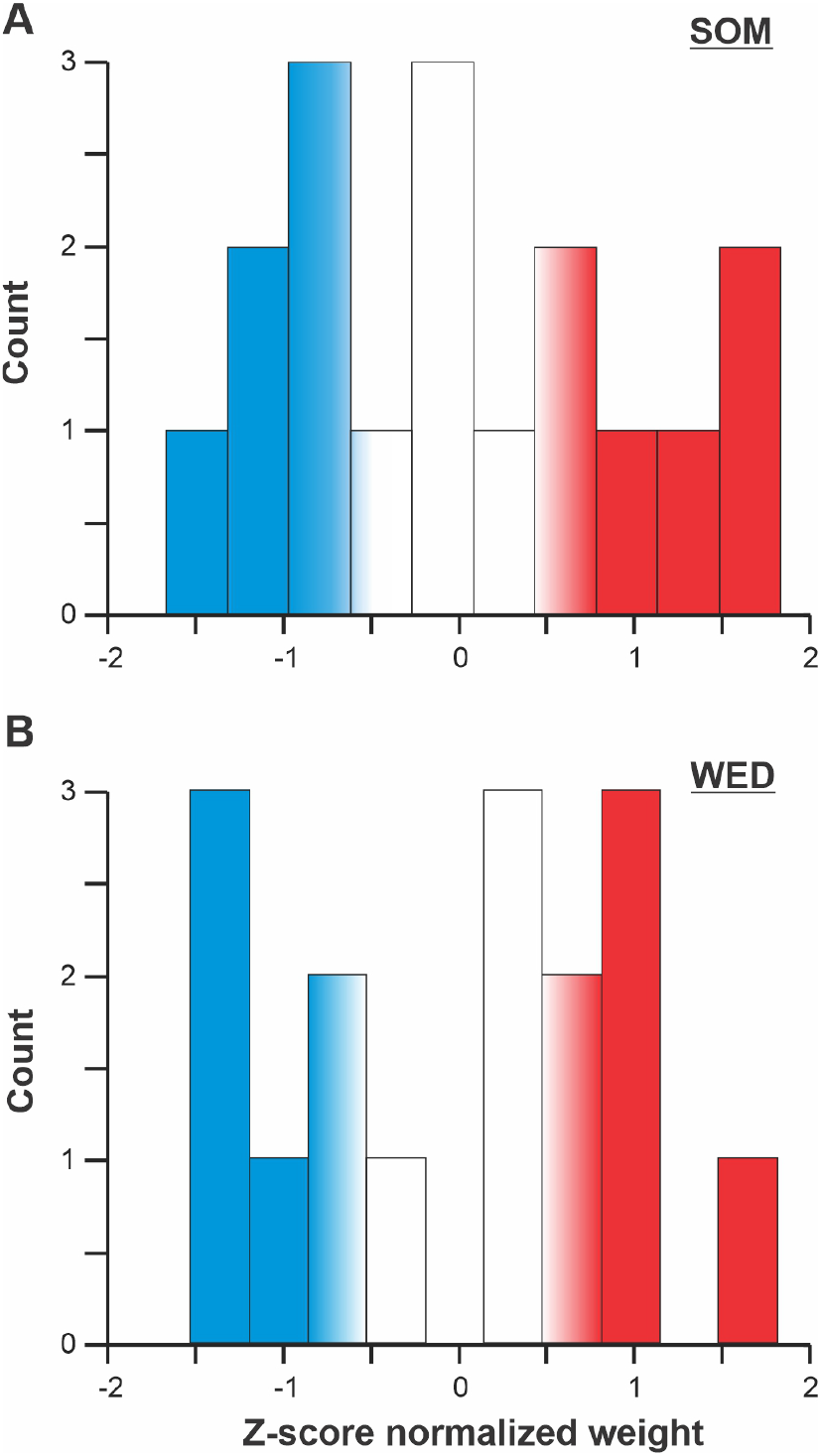
Distribution of weights for each of the 16 neurons in one analysis of discrimination for a pair of odor stimuli. **A.** Weights extracted from the trained SOM neural net performing the discrimination task. **B.** Weights for the WED discrimination analysis of the same pair of stimuli. The weights are z-score normalized for comparison. We see that efficient discrimination required ~ 6 neurons to be strongly weighted (highlighted in red; 8-9 neurons were weighted more than the average). This secondary analysis could be useful as a prediction of the connectivity patterns that could present in the biological system (e.g. how many neurons converged onto a single decoding postsynaptic cell).

**Supplementary Information Figure 5:**
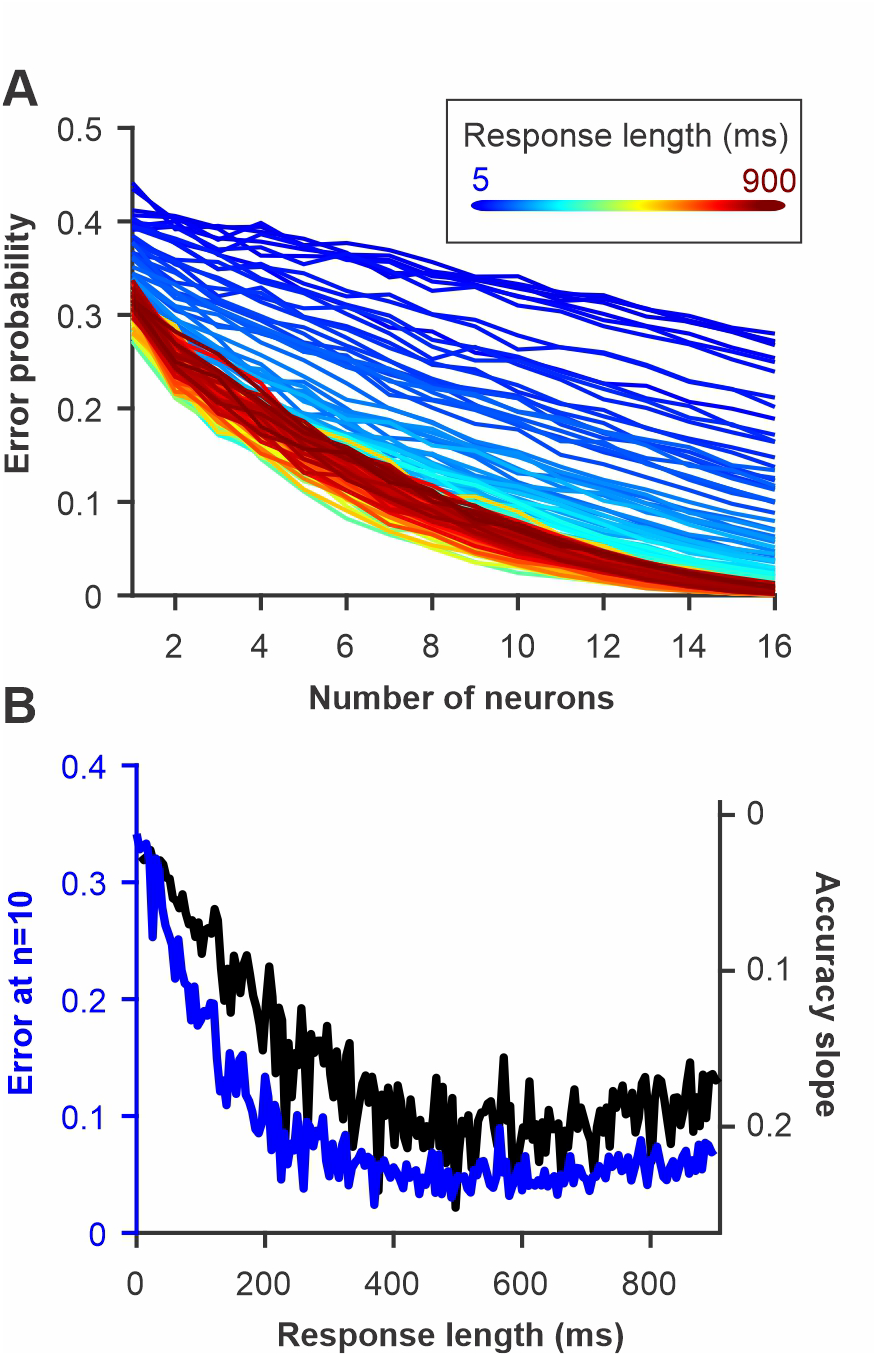
Discrimination accuracy for pairs of olfactory stimuli as increasing lengths of the neural responses after stimulus onset are included in the analysis. **A.** Error probability as a function of number of neurons included in the analysis and duration of the response. **B.** Discrimination performance quantified by the error rate for a population of 10 neurons (blue) or as the slope of the curves in A (see Fig 6). We see that discrimination accuracy should increase quickly in the first 200 ms of the response but then plateaus. These estimates can be related to the behavioral performance of the animal. For behavioral response that happen quickly after stimulus onset, neural responses pattern should contain enough information in the initial portion of their responses to support accurate discrimination. Interestingly, it has been shown that moths can perform accurate behavioral discrimination within a few hundred ms of the odor presentation for these type of stimuli.

**Supplementary Information Figure 6:**
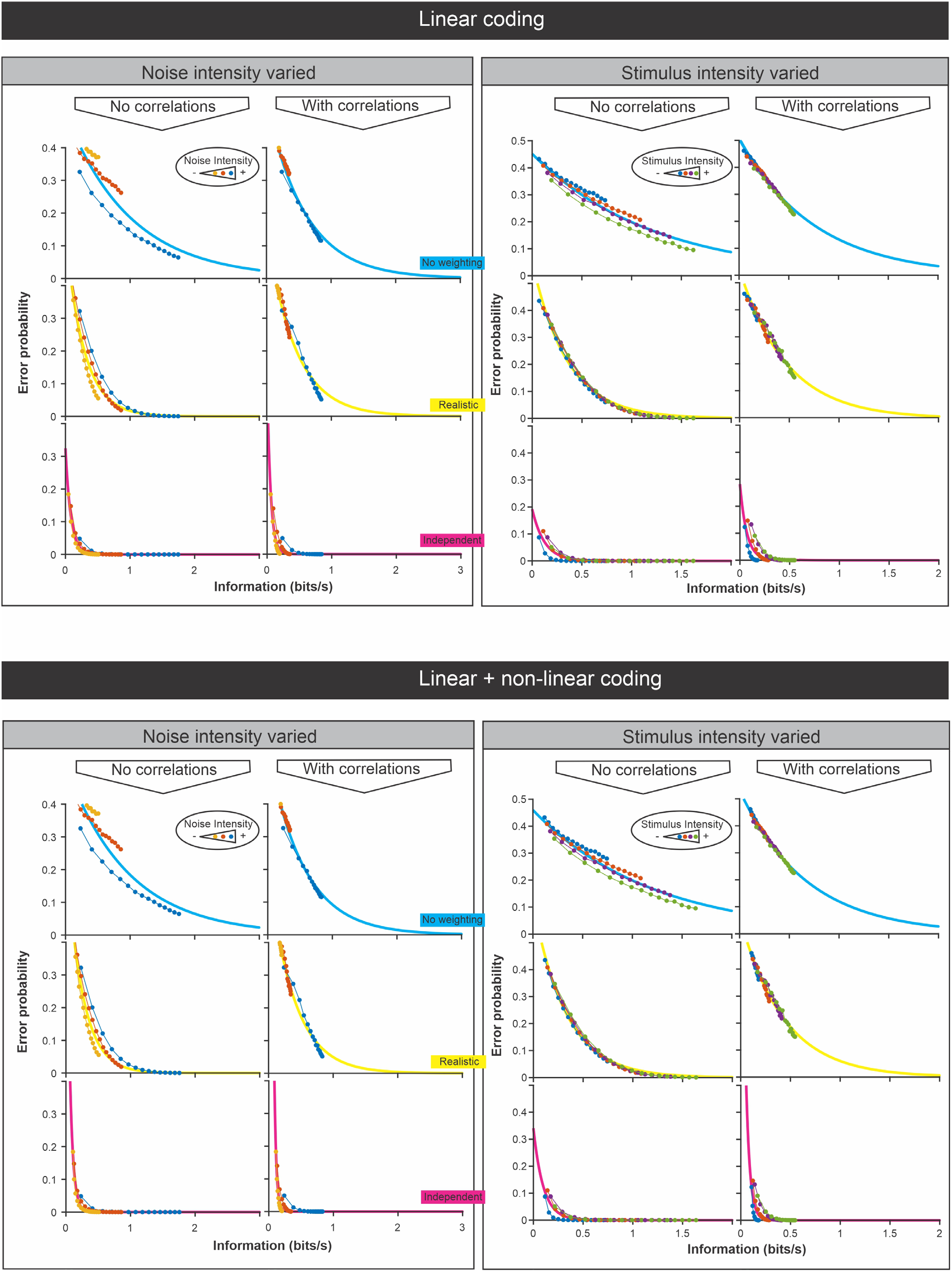
Comparison of discrimination error probability and information content of LIF model responses to frozen white noise stimuli. This figure complements Fig 7 of the main text. We performed the analysis using response-stimulus coherence to estimate information, which give a lower-bound estimate of the information content because it considers only the information encoded linearly. We also performed the analysis using response-response coherence which provides an upper-bound estimate of information content that includes both linear and non-linear coding (see Borst and Theunissen 1999 for more details). Our LIF model was adjusted to be in a linear regime therefor there is no difference between the two sets of plots (upper half vs bottom half of the figure). To change the signal to noise ratio we changed the amount of noise while keeping the strength of the signal constant (left half of this figure; similar to Fig 7 although here the overall signal strength was lower than in Fig 7). Alternatively, we changed the stimulus strength keeping the noise constant (right half of this figure). The analysis was performed with noise correlation present of not (see Figure 8 for more details). Each panel presents the analysis with the three different weighting methods (no weighting in blue, realistic weighting in yellow and optimized weighting in magenta) that are used in Fig 6–8. We fitted exponential curves in each plot to show the high correlation between information content and discrimination accuracy. We note that this relationship changes with the discrimination decoding methods as some methods can leverage better the most informative parts of the responses and thus lead to better discrimination although the information content doesn’t change. The relationship can also be affected but the characteristics of the stimulus as shown by the fact that results for different signal-to noise ratios follow slightly different curves in some plots (compare for example the 3 curves in the “realistic” analysis when the noise intensity is varied and no noise correlations are present).

## Notes

### Competing Interest Statement

The authors have declared no competing interest.

### Summary of Updates

In this revision, we added details to the methods and several supplementary figures

